# Information processing in biological molecular machines

**DOI:** 10.1101/2020.01.24.918367

**Authors:** M. Kurzynski, P. Chelminiak

## Abstract

Biological molecular machines are enzymes that simultaneously catalyze two processes, one donating free energy and second accepting it. Recent studies show that most native protein enzymes have a rich stochastic dynamics of conformational transitions which often manifests in fluctuating rates of the catalyzed processes and the presence of short-term memory resulting from the preference of certain conformations. For arbitrarily complex stochastic dynamics of protein machines, we proved the generalized fluctuation theorem predicting the possibility of reducing free energy dissipation at the expense of creating some information stored in memory. That this may be the case has been shown by interpreting results of computer simulations for a complex model network of stochastic transitions. The subject of the analysis was the time course of the catalyzed processes expressed by sequences of jumps at random moments of time. Since similar signals can be registered in the observation of real systems, all theses of the paper are open to experimental verification.

**STATEMENT OF SIGNIFICANCE:** The transient utilization of memory for storing information turns out to be crucial for the movement of protein motors and the reason for most protein machines to operate as dimers or higher organized assemblies. From a broader physical point of view, the division of free energy into the operation and organization energies is worth emphasizing. Information can be assigned a physical meaning of a change in the value of both these functions of state.

## 1 Introduction

The basic task of statistical physics is to combine the dynamics of the microstates of a studied system with the dynamics of the macrostates. The transition from microstates to macrostates is the result of averaging over a sufficiently long period of time [1]. In the statistical physics of simple systems, there are no intermediate levels of organization between microscopic mechanics and macroscopic thermodynamics. Living matter is, however, a complex system with the entire hierarchy of organization levels [2, 3]. The new achievements at the turn of the century allowed us to understand more accurately the nature of the micro and macrostates of the biological molecular machines, which belong to the level currently being referred to as nanoscopic.

First, the tertiary structure of the enzymatic proteins, formerly identified with one or at most several conformational states, has been extended to the whole network of conformational substates [4, 5, 6, 7, 8, 9, 10, 11, 12, 13]. The Markov stochastic process on such a network is to be considered as the starting microscopic dynamics for the statistical treatment. The rule are fluctuating rates of the catalyzed reactions (the “dynamic disorder”) [8, 9, 14], which means realization of them in various randomly selected ways. As a symbol of the progress made recently, the molecular dynamics study of conformational transitions in the native phosphoglycerate kinase could be quoted, in which a network of 530 nodes was found in the long 17 *μ*s simulation [15]. This network seems to display a transition from the fractal to small-world organization [16, 17]. On such networks, modeled by scale-free fractal trees [18] extended by long-range shortcuts, the random choice of the course of catalyzed processes is quite natural [19].

Secondly, new methods of stochastic thermodynamics have been applied to the description of the non-equilibrium behavior of single nanoobjects in a finite time perspective. Work, free energy dissipation and heat in nanoscopic systems are random variables and their fluctuations, proceeding forward and backward in time, proved to be related to each other by the fluctuation theorem [20, 21, 22, 23]. For nanoscopic machines, the concept of information processing can be defined and the relationships between information and entropy production leads to the generalized fluctuation theorem [24, 25, 26, 27, 28, 29, 30, 31, 32]. It strengthened an almost universal consensus regarding the view on the operation of Maxwell’s demon consistent with thermodynamics [33, 34, 35, 36, 37]. Thus, the demon must have a memory and in order to use fluctuations to do the work, it reduces entropy at the expense of gathering the necessary information on fluctuations in this memory. The information must be erased sooner or later, and for this the same or greater work must be used.

The use of the language of stochastic thermodynamics to describe the operation of biological molecular machines requires precise definition of the concept of the nanoscopic machine. In the thermodynamic context, a machine means any physical system that allows two other systems to perform work one on another. Work can be done by mechanical, electrical, chemical, thermal or still some other forces. The forces are defined as the differences of the respective potentials [38]. Because the biological molecular machines operate using thermal fluctuations, just like chemical reactions, we treat them as chemo-chemical machines [39]. The protein chemo-chemical machines are to be considered as enzymes, that simultaneously catalyze two effectively unimolecular chemical reactions: the energy-donating input reaction R_1_ → P_1_ and the energy-accepting output reaction R_2_ → P_2_, see Fig. 1 A. Also pumps and molecular motors can be treated in the same way, see Figs. 1 B and C.

**Figure 1:**
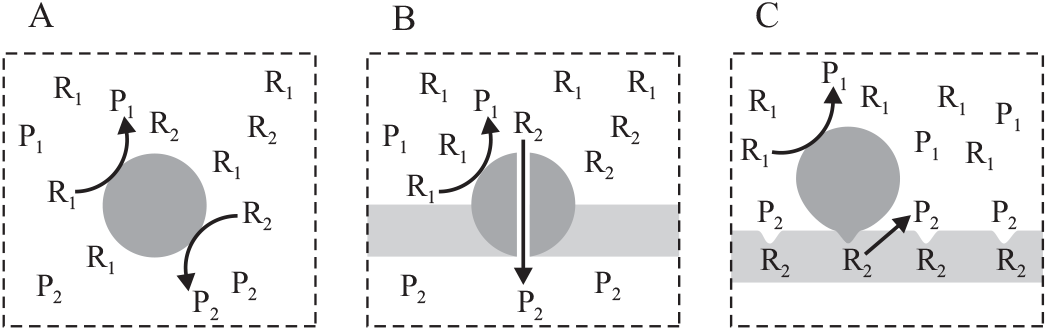
Schematic picture of the three types of the biological molecular machines. (A) Enzymes that simultaneously catalyze two reactions. (B) Pumps placed in a biological membrane – the molecules present on either side of the membrane can be considered to occupy different chemical states. (C) Motors moving along a track – the external load influences the free energy of motor binding, which can be considered as a change in the probability of finding a particular track’s binding site, hence its effective concentration [39, 40].

The chemo-chemical machine we consider consists of a single enzyme macromolecule, surrounded by a solution of its substrates and products (Fig. 1). The whole is an open non-equilibrium system which operates stochastically under the influence of forces controlled by the concentrations of reactants. Under specified relations between the concentrations of all reactants [39], the non-equilibrium molar concentrations of product molecules [P_1_] and [P_2_], related to enzyme total concentration [E], are the two dimensionless thermodynamic variables *X*_1_ and *X*_2_ determining the state of the system. These, together with the conjugate thermodynamic forces, i.e. chemical affinities (the differences in chemical potentials) *A*_1_ and *A*_2_, determine work performed on and by the machine, respectively. Assuming that the molecule solution is perfect, the formal definitions have the form [39, 41]:

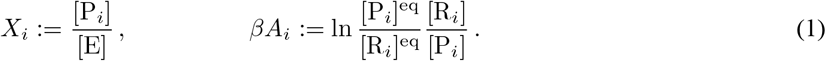

*β* is the reciprocal of the thermal energy *k*_B_*T* and the superscript eq denotes the equilibrium concentrations.

The observable action of the machine is its turnovers under the influence of forces *A*_1_ and *A*_2_ measured by the number of molecules P_1_ and P_2_ created in a unit of time. Establishing the forces *A*_1_ and *A*_2_ and the variables *X*_1_ and *X*_2_ defining them is formally equivalent to holding the machine in the steady-state conditions due to imagined constraints that control the number of incoming and outgoing reacting molecules, what is symbolically illustrated by a frame of a broken line in Fig. 1. It should be stressed that the entire system inside the frame represents the machine: it is these constraints that determine the values of the thermodynamic variables and the conjugate forces acting on the machine.

The biological molecular machines operate at a constant temperature. Under isothermal conditions, the internal energy is uniquely divided into free energy, the component that can be turned into work, and entropy multiplied by temperature (after Helmholtz, we call it bound energy), the component that can be turned into heat [38]. Both thermodynamic quantities can make sense in the non-equilibrium state if the latter is treated as a partial equilibrium state [39]. Free energy can be irreversibly transformed into bound energy in the process of internal entropy production which, in the energy-related context, means free energy dissipation. In view of such internal energy division, the protein molecular machines are referred to as free energy transducers [41]. The energy processing pathways in a stationary isothermal machine are shown in Fig. 2 A, where the role of all the physical quantities being in use is also indicated. *X*_*i*_ denotes the input (*i* = 1) or the output (*i* = 2) thermodynamic variable, *A*_i_ is the conjugate thermodynamic force and the time derivative, *J*_i_ = d*X*_*i*_/d*t*, is the corresponding flux. *T* is the temperature and *S* is the entropy. To clearly specify the degree of coupling of the fluxes, *ϵ* := *J*_2_/*J*_1_, it is important that variables *X*_1_ and *X*_2_ be dimensionless as in Eq. 1. Corresponding forces *A*_1_ and *A*_2_ are then also dimensionless, if only multiplied by the reversal of the thermal energy *β* = (*k*_B_*T*)^−1^. We assume *J*_1_, *J*_2_, *A*_1_ > 0 and *A*_2_ < 0 throughout this paper but −*A*_2_ ≤ *A*_1_, i.e., variables *X*_1_ and *X*_2_ are defined such that the machine does not work as a gear.

**Figure 2:**
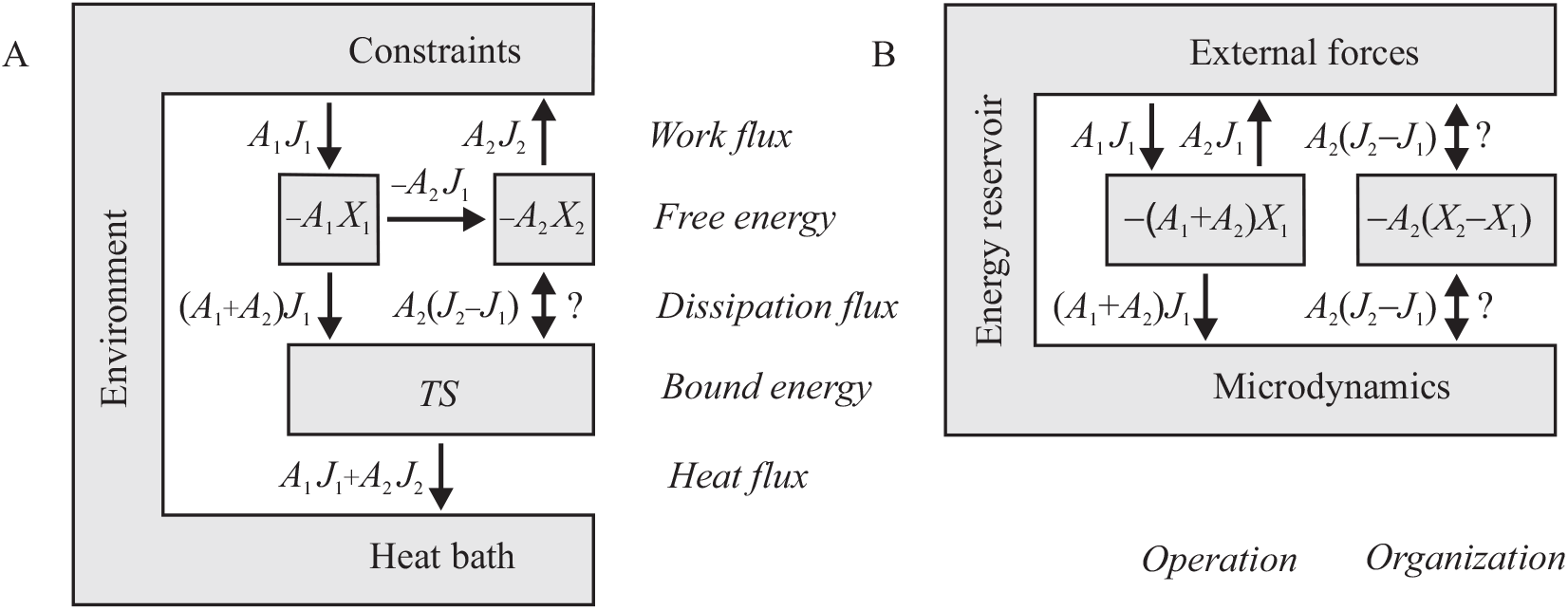
Energy processing in the stationary isothermal machine. The constraints keep stationary values of thermodynamic variables *X*_1_ and *X*_2_ fixed. (A) Division of the machine’s internal energy into free energy *F* = −*A*_1_*X*_1_ − *A*_2_*X*_2_ and bound energy *T S*. The directions of the energy fluxes shown are for *J*_1_, *J*_2_, *A*_1_ > 0 and *A*_2_ < 0. The direction of flux *A*_2_(*J*_2_ − *J*_1_), marked with a question mark, is a subject of discussion in this paper. (B) The alternative view described in the text. Only free energy, observable macroscopically, is specified. The bound energy, determined by unknown microdynamics, is considered, along with the heat bath and imagined constraints, as the energy reservoir.

In the steady state, the resultant work flux (the processed power) *A*_1_*J*_1_ + *A*_2_*J*_2_ equals the heat flux and both equal the dissipation flux (the rate of internal entropy production multiplied by the temperature), which, according to the second law of thermodynamics, must be non-negative. However, it consists of two components. The first component, (*A*_1_ + *A*_2_)*J*_1_, achieved when the fluxes are tightly coupled, *J*_2_ = *J*_1_, must also be, according to the same law, non-negative, but the sign of the complement *A*_2_(*J*_2_ − *J*_1_) is open to discussion. In the macroscopic machines, the latter is also non-negative and has the obvious interpretation of a frictional slippage in the case of mechanical machines, a short-circuit in the case of electrical machines, or a leakage in the case of pumps [39]. However, because of non-vanishing correlations [24, 25, 26, 27, 28, 29, 30, 31, 32], entropy *S* in the nanoscopic machines is not additive as in the macroscopic machines, which could cause a change the sign of *A*_2_(*J*_2_ − *J*_1_) to negative, i.e., for *A*_2_ < 0, to *J*_2_ ≥ *J*_1_.

From the point of view of the output force *A*_2_, subsystem 1 carries out work on subsystem 2 while subsystem 2 carries out work on the environment. Jointly, the flux of the resultant work (the resultant power) *A*_2_(*J*_2_ − *J*_1_) is driven by the force *A*_2_. The complement to *A*_1_*J*_1_ + *A*_2_*J*_2_ is flux (*A*_1_ + *A*_2_)*J*_1_ driven by the force *A*_1_ + *A*_2_. Consequently, the free energy processing from Fig. 2 A can be alternatively presented as in Fig. 2 B, with the free energy transduction path absent. Here, two interacting subsystems of the machine characterized by variables *X*_1_ and *X*_2_ have been replaced by two non-interacting subsystems characterized by variable *X*_1_ and the difference *X*_2_ − *X*_1_. The first describes the perfect operation of the machine while the other its deteriorating or improving organization. Therefore, we will call *X*_1_ the machine operational variable, and *X*_2_ − *X*_1_ the machine organizational variable. As a result of interaction with microdynamics, both newly defined subsystems, although non-interacting directly, still remain statistically correlated.

The purpose of this paper is to answer the question of whether the particular stochastic dynamics of the biological molecular machines allows them to act as Maxwell’s demons. This would help to understand the hard-to-interpret behavior of single-headed biological motors: myosin II [42], myosin V [43], kinesin 3 [44], and both flagellar [45] and cytoplasmic [46] dynein, which can take more than one step along their track per ATP or GTP molecule hydrolyzed. This means that the ratio *ϵ* = *J*_2_/*J*_1_ is higher than unity, so the sign of dissipation flux *A*_2_(*J*_2_ − *J*_1_)A in Fig. 2 B is negative.

Basing on recent investigations, we assume that the stochastic dynamics of the enzymatic machine is characterized by transient memory and the possibility to realize the catalyzed process in various randomly selected ways. For such dynamics we calculate both the free energy dissipation and information, and show that the first can change the sign being partially compensated by the second. Because available experimental data on the actual conformational transition networks in native proteins is still very poor, we verify the theoretical results only by computer simulations for a model network. Our conclusions are based on analysis of the simulated time course of the catalyzed processes expressed by sequences of jumps in the values of observed variables at random moments of time. Since similar signals can be registered in the experiments [42, 43, 44, 45, 46], all theses of the paper are open for experimental verification. Thus, besides presentation of new theoretical concepts, the paper can be also treated as an invitation to experimentalists to perform similar analysis on real systems.

## 2 Methods

### 2.1 Bipartite dynamics of biological molecular machines

In the language of conventional chemical kinetics, the operation of a chemo-chemical machine is explained by the scheme in Fig. 3 A: the energy-donating reaction R_1_ → P_1_ forces the direction of the energy-accepting reaction R_2_ → P_2_, though the nonequilibrium concentration values [R_2_] and [P_2_] would prefer the opposite direction [39, 41]. We assumed that the binding of reactants precedes their detaching but this order may be reversed as shown in Fig. 3 B. In both cases, for the sake of simplicity, we do not take into account details of the binding and detaching reactions involving spatial diffusion of reactants and molecular recognition [39, 47].

**Figure 3:**
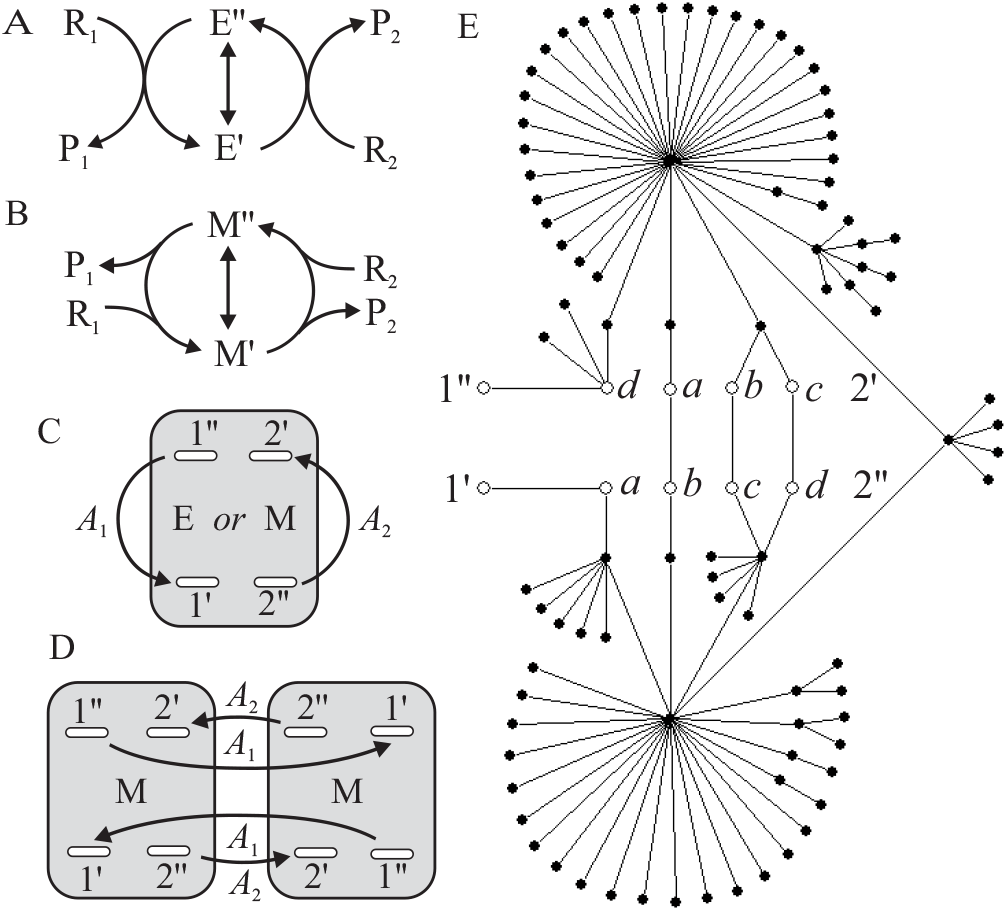
Dynamics of the enzymatic chemo-chemical machine. (A) and (B) Schemes of conventional chemical kinetics that take into account only two states E″ and E′ of the free enzyme or the two states M″ and M′ of the enzyme-reagents complex. (C) The generalized scheme [19, 47]. The gray box represents a one-component network of transitions between conformational substates of the enzyme or the enzyme-reactants complex native state. All these transitions satisfy the detailed balance condition. A system of single or multiple (ovals) pairs of conformational substates (the “gates”) (1″, 1′) and (2″, 2′) has been distinguished, between which input reaction R_1_ → P_1_ and output reaction R_2_ → P_2_ cause, respectively, additional transitions that break the detailed balance. All the reactions are reversible; the arrows indicate the directions assumed to be forward. (D) Bipartite structure of the formally doubled network described in the text. (E) Sample implementation of the 100-node network, constructed following the algorithm described in the text. For simplicity, a single input gate is assumed. A single output gate chosen for the simulations is created by the pair of transition states (2″*a*, 2′*d*). The four pairs of transition states (2″*a*, 2′*a*), (2″*b*, 2′*b*), (2″*c*, 2′*c*) and (2″*d*, 2′*d*) lying tendentiously one after another, create alternative fourfold output gate.

Only two states E′ and E″ of the free enzyme are distinguished in the scheme in Fig. 3 A and only two states M′ and M″ of the enzyme-reagents complex are distinguished in the scheme in Fig. 3 B. Only one non-reactive transition between them is assumed. However, as mentioned in the Introduction, the nanoscopic kinetics of biological machines is described by the whole network of conformational substates. In this network, with transitions that obey the detailed balance condition, a system of pairs of nodes (the “gates”) is distinguished, between which the input and output chemical reactions force additional transitions that break the detailed balance in a degree dependent on the values of *A*_1_ and *A*_2_ [19, 39, 47], see Fig. 3 C. The rule is the presence of many gates allowing the choice of the way of implementing reactions [8, 9].

The scheme in Fig. 3 C determines fluxes *J*_1_ and *J*_2_ in a statistical ensemble of many molecular machines, but does not predict the result of a nanoscopic observer who counts the molecules P_1_ and P_2_ successively created or destroyed by a single machine, in particular finding its location on the track. Including the observer’s memory in this scheme would mean non-Markovianity of the process, but this can be avoided by formal doubling the network [30, 31, 32] and replacing the scheme in Fig. 3 C with that presented in Fig. 3 D. Now, the state of the machine is determined not only by the network node *m* but also by the gate *g* through which it has recently passed. In other words, the space of states of the machine is a subset of the Cartesian product **G** × **M**, where **M** is a set of all nodes (we consider they represent the states of the enzyme-substrates complex) and **G** is a set of nodes that are simultaneously the gates transition states.

All the transitions within the doubled network in Fig. 3 D can be divided into internal transitions of the form (*g, m*) → (*g, m*′) (the index of the recent crossed gate *g* is established) and external transitions of the form (*g, m*) → (*g*′, *m*) (the passage through a specific gate can be uniquely identified with the corresponding final conformational substate, *m* = *g*′). This means that the doubled network has a bipartite structure, which is a sufficient condition to clearly define the concept of information processing [26, 27, 28, 29, 30, 31, 32]. The internal transitions are hidden for the observer who only sees the external transitions. Averaging the stochastic trajectory in the space of states from Fig. 3 D over a certain period of time *t* determines the resultant number of P_1_ and P_2_ molecules created at time *t*, i.e., the random values of fluxes *j*_1_(*t*) and *j*_2_(*t*) tending, for *t* → ∞, to *J*_1_ and *J*_2_.

### 2.2 Specification of the computer model

Because available experimental data on the actual conformational transition networks in native proteins are still very poor, we constructed a model network to apply the theory for interpreting results of computer simulations. As mentioned in the Introduction, this network should be scale-free and display a transition from the fractal to small-world organization [15, 16, 17, 18, 19]. A sample network of 100 nodes with such property is depicted in Fig. 3 E. The algorithm of constructing the stochastic scale-free fractal trees was taken from Goh et al. [18]. Shortcuts, though more sparsely distributed, were considered after Rozenfeld, Song and Makse [16]. Here, we randomly chose three shortcuts from the set of all the pairs of nodes distanced by six links. The network of 100 nodes in Fig. 3 E is too small to determine its scaling properties, but a similar procedure of construction applied to 10^5^ nodes results in a scale-free network, which is fractal on a small length-scale and a small world on a large length-scale.

To provide the network with stochastic dynamics, we assume the probability of changing a node to any of its neighbors to be the same in each random walking step [19, 48]. Then, following the detailed balance condition, the equilibrium free energy of a given node is inversely proportional to the number of its links (the node degree). The most stable nodes are the hubs. The assumed dynamics is independent of temperature, and therefore not very realistic, but this problem is not the subject of our research here. For given node *l*, the transition probability to any of the neighboring nodes per random walking step is one over the number of links and equals 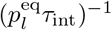, where 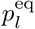 is the equilibrium occupation probability of the given node and *τ*_int_ is the mean time to repeat a chosen internal transition, counted in the random walking steps. This time is determined by the doubled number of links minus one [48], *τ*_int_ = 2(100 + 3 − 1) = 204 random walking steps for the 100 node tree network with 3 shortcuts assumed.

In the scheme in Fig. 3 E, let us note two hubs, the protein substates of the lowest free energy, e.g. “open” and “closed”, or “bent” and “straight”, usually the only ones occupied sufficiently high to be observable under equilibrium conditions. The dream of structural biologists is to find the largest possible number of such distinguished states for actual protein machines. The proposed kind of stochastic dynamics combines in fact a chemical description with a mechanical one [39] but let us remember that in contrast to macroscopic mechanical machines, the transitions between such states in nanomachines are always stochastic.

The external transition probability competes with the probability of possible internal transitions and, for the forward transition, equals 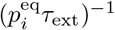 per random walking step, 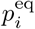 denoting the equilibrium occupation probability of the initial input or output node (*i* = 1, 2, respectively). The corresponding backward external transition probability is modified by detailed balance breaking factors exp(−*βA*_i_). The mean time of forward external transition *τ*_ext_ is related to stationary concentration [P_i_]. We assume *τ*_ext_ = 20 random walking steps, one order of the magnitude shorter than the mean time of internal transition *τ*_int_, which means that the whole process is controlled by the internal dynamics of the system. Identifying a time unit with a computer step still requires establishing the probabilities of staying, different on network and gate nodes [48].

## 3 Results and Discussion

### 3.1 Generalized fluctuation theorem for biological molecular machines

The bipartite structure of the stochastic dynamics of biological molecular machines presented in Fig. 3 D makes possible treating transitions under the influence of external forces as a distinguished subset in which partial entropy production takes place, specified by the fluctuation theorem in the form of the Jarzynski equality [31, 32]:

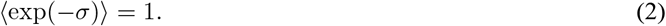

The stochastic dimensionless entropy production *σ* in the external transitions, multiplied by the thermal energy *k*_B_*T* = *β*^−1^, determines free energy dissipation for a period of time *t* in the stationary energy processing presented in Fig. 2, hence

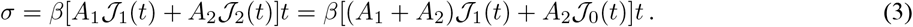

Here, the random variables are the net fluxes 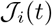 averaged over the time interval *t*, *i* = 1, 2 or *i* = 1, 0. We wrote two alternative divisions of the dissipation flux discussed in Fig. 2 and we defined

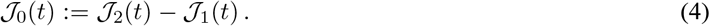

The detailed fluctuation theorem corresponding to the integral fluctuation theorem 2 with the second division 3 is, in the logarithmic form,

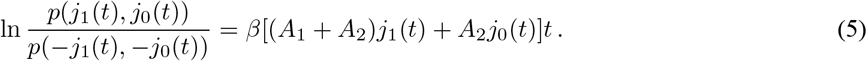

*j*_*i*_(*t*) in 5, *i* = 1, 0, denote particular values of random fluxes 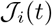 in 3. *p* is the joint probability distribution function of these fluxes and the average in 2 is performed over this distribution. By definition, the stationary averages of the random fluxes are time independent, 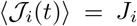 for any *t*, but the value of the higher moments of distribution *p*(*j*_1_(*t*), *j*_0_(*t*)) increases with the shortening of time *t*. The convexity of the exponential function provides the second law of thermodynamics

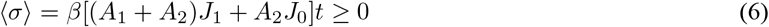

to be a consequence of the Jarzynski equality 2.

Equation 5 is identical to the fluctuation theorem for the stationary fluxes earlier introduced by Gaspard [49] who based it on much more complex considerations, but the condition on *t* can be more precisely determined here. The chosen time *t* must be long enough for the considered ensemble to comprise only stationary fluxes, but finite to observe any fluctuations. As all stationary fluxes in the ensemble are statistically independent, the probability distribution function *p*(*j*_1_(*t*), *j*_0_(*t*)) is the two-dimensional Gaussian [50]

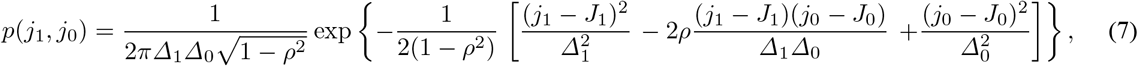

where the averages 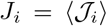, *i* = 1, 0, and the corresponding variances (the squares of the standard deviations) 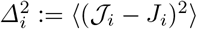 specify the Gaussian marginals for the individual fluxes, and *ρ* is the correlation coefficient:

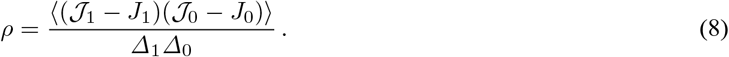

Here and further on, for the sake of brevity, we omit argument *t* specifying all the fluxes. The requirement for the Gaussian 7 to satisfy the detailed fluctuation theorem 5 leads to a system of two quadratic equations that link standard deviations Δ_1_ and Δ_0_ with averages *J*_1_ and *J*_0_:

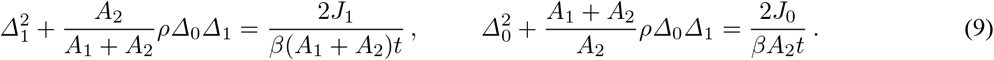

From 9, it follows that standard deviations *Δ*_1_ and *Δ*_0_ are inversely proportional to the square root of *t*. For the lack of correlations, *ρ* = 0, the solutions to 9 reconstruct our earlier result [48].

Eqs. 9 allow to calculate the ratios of marginals *p*(*j*_1_)*/p*(−*j*_1_) and *p*(*j*_0_)*/p*(−*j*_0_) for fluxes 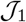 and 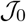 separately, from which it follows that the detailed and integral fluctuation theorem takes the form

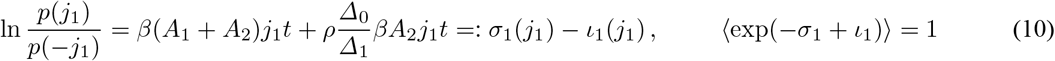

for the flux 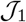 and

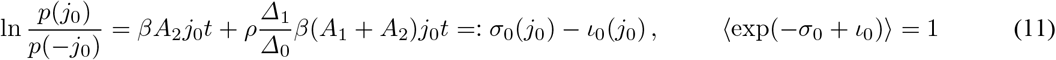

for the flux 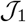. The averages are performed over the one-dimensional distributions, either *p*(*j*_1_) or *p*(*j*_0_). In addition to the functions *σ*_1_ and *σ*_0_ that describe separate entropy productions, we have obtained additional functions *ι*_1_ and *ι*_0_. As Δ_1_ and Δ_0_ are positive and forces *β*(*A*_1_ + *A*_2_) and *βA*_2_ are of the opposite signs, components *σ*_i_ and *ι*_i_ substracts when *ρ* is positive and add up when *ρ* is negative.

The interpretation of functions *ι*_1_ and *ι*_0_ is not obvious. Because there are no random variables that specify the complements of fluxes 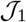 and 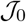 treated separately, the expressions for *ι*_1_ and *ι*_0_ provided by Eqs. 10 and 11 cannot be rewritten using the concept of Shannon’s mutual information [26, 27, 28]. Only the sum of both functions *ι*_1_ and *ι*_0_ is related to mutual information between macrovariables *J*_1_ and *J*_2_, as from Eqs. 5, 10 and 11, the relationship

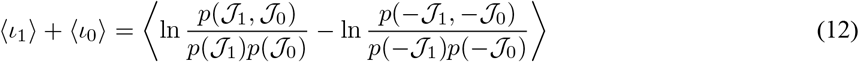

results. Let us emphasize that because *ι*_1_ is the function of only *j*_1_, and *ι*_0_ is the function of only *j*_0_, the value of the difference of the logarithms in 12 remains the same when averaged over the correlated distribution *p*(*j*_1_, *j*_0_) or the product *p*(*j*_1_)*p*(*j*_0_). Of course, this is not true for the separate components of the logarithms difference in 12.

The functions *ι*_1_ and *ι*_0_, however, actually represent the information exchanged between microdynamics and macrodynamics if only the former is treated as an information reservoir [28, 29, 30, 31, 32, 37, 51]. We mentioned in the Introduction that the observable action of the machine is its turnovers measured by the net number of the created molecules P_1_ and P_2_. Hence, the signal sent by the machine is in the form of two sequences of bits, or rather signs, e.g. …, +, +, −, +, −, +, +, +, −, −, +, …, that describe successive transitions forward or back through the input or output gates, respectively.

Because Eq. 12 is symmetric with respect to the replacement of 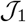 with 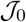, it does not describe any directed information transfer between both macrodynamics. This means that in addition to the law of free energy conservation in the steady state in each of the subsystems *i* = 1, 0, there is an additional law of conservation stating the equality of information transmitted from microdynamics to macrodynamics with information transmitted from macrodynamics to the environment as a signal of random passage of product molecules through the constraints. Indeed, with the help of Eqs. 9, Eqs. 10 and 11 can be rewritten directly as

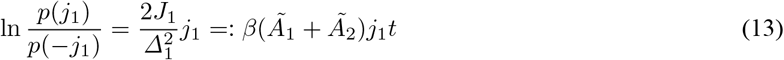

for the flux 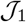 and

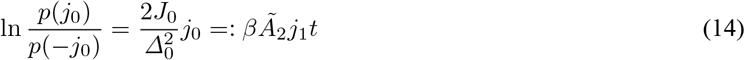

for the flux 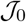. Forces 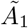 and 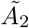 defined above include, in addition to the forces *A*_1_ and *A*_2_, corrections to the signals of random passages of the product molecules through the constraints. These signals play the role of additional constant chemical forces [32] acting on each of the macrodynamics separately. In such approach, the remained macrodynamics together with microdynamics are jointly treated as hidden [52] what is presented in Fig. 4 A.

**Figure 4:**
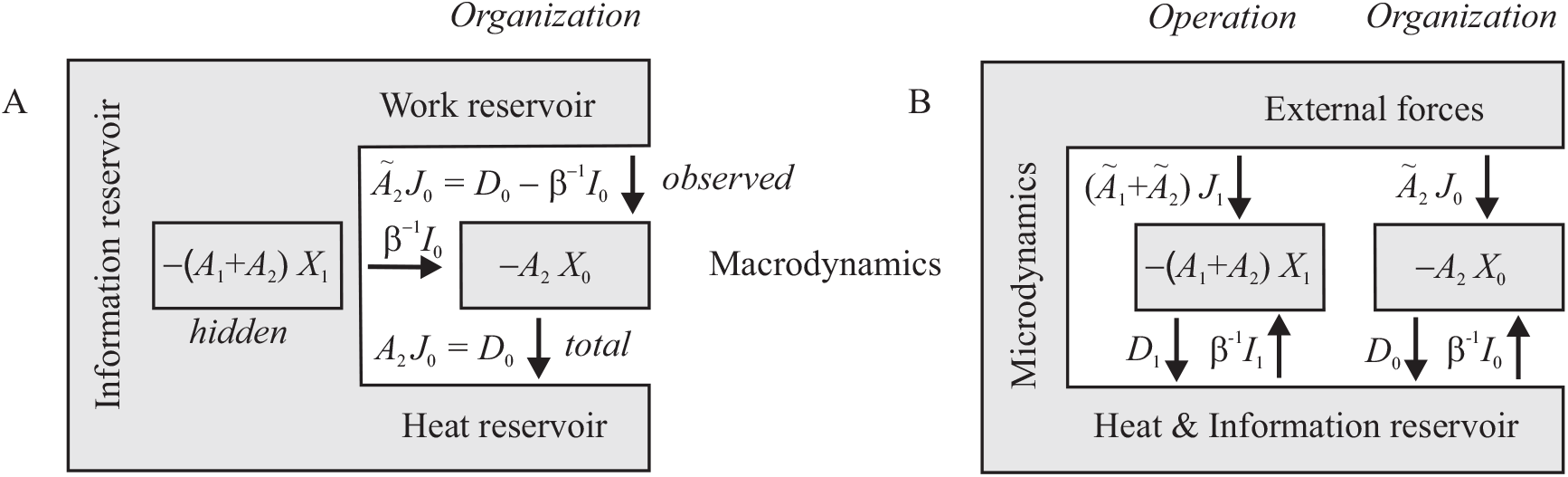
(A) For each separately observed macrodynamics, the second macrodynamics is hidden and can be treated together with microdynamics as an information reservoir. The case of an organizational variable is shown. (B) General scheme of communication between macrodynamics, which determines free energy, and microdynamics. 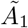 and 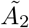 denote the effective forces that take additionally into account the signals coming from the environment.

A comparison of two equivalent formulations of the fluctuation theorem for separate fluxes, 10 and 11 with 13 and 14, leads to expressions for stochastic informations *ι*_1_ and *ι*_2_ dependent only on standard deviations Δ_1_ and Δ_2_ but not on the correlation coefficient *ρ*. Only four parameters *J*_i_ and Δ_i_, *i* = 1, 0, that characterize the Gaussian distribution functions of the separate fluxes, are sufficient to specify the four averages: dissipation multiplied by reciprocal thermal energy *βD*_i_ := 〈*σ*_i_〉 and information *I*_i_ := 〈*ι*_i_〉 for *i* = 1 and 0, respectively. From integral fluctuation theorems (10) and (11), it follows that they satisfy the generalized second laws of thermodynamics:

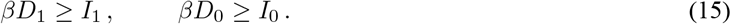

Fig. 4 B presents a general communication between macrodynamics, characterized by free energy, and microdynamics combined with the environment, including both dissipation and exchange of information. *D*_1_ is always positive, but if *I*_0_ is negative, also *D*_0_ can be negative, which means the organizational subsystem may behave like Maxwell’s demon.

### 3.2 Application of analytical results to the model system

Because contributions from dissipation *D*_1_ and *D*_2_ depend only on the values of stationary fluxes *J*_1_ and *J*_2_, they can easily be determined for fixed values of forces *A*_1_ and *A*_2_ [19]. More detailed statistics are needed to determine contributions from information *I*_1_ and *I*_2_. The general theory does not provide for both the *D*_0_ sign and the *I*_1_ and *I*_0_ signs. To check if the organizational subsystem of the biological molecular machine can actually behave like Maxwell’s demon, we performed computer simulations of random walk on the network constructed following the algorithm described in the Methods and shown in Fig. 3 E.

We assumed *βA*_1_ = 1 and a few smaller, negative values of *βA*_2_. The result of each simulation was the time courses of the net numbers *x*_1_ and *x*_2_ of transitions through the input and output gates, respectively. Exemplary courses are shown in Fig. 5 A. The thermodynamic description of the nanosystem requires averaging over a finite observation period. To get statistical ensembles of the stationary fluxes, we divided long stochastic trajectories of 10^10^ computer steps into segments of equal lengths *t*. For the fluxes *j*_i_ to be statistically independent, the selected time *t* must have been longer than the complete cycle of free energy transduction. Although the microscopic dynamics is a Markov process, the time courses *x*_1_ and *x*_2_ are not Markov processes but continuous time random walks [50, 53]. By analyzing their dwell time distributions, we found the reasonable time of averaging to be *t* = 1000 computer steps. The result of averaging is pairs of fluxes *j*_i_(*t*) = Δ*x*_i_(*t*)*/t*, *i* = 1, 2, which is also shown in Fig. 5 A.

**Figure 5:**
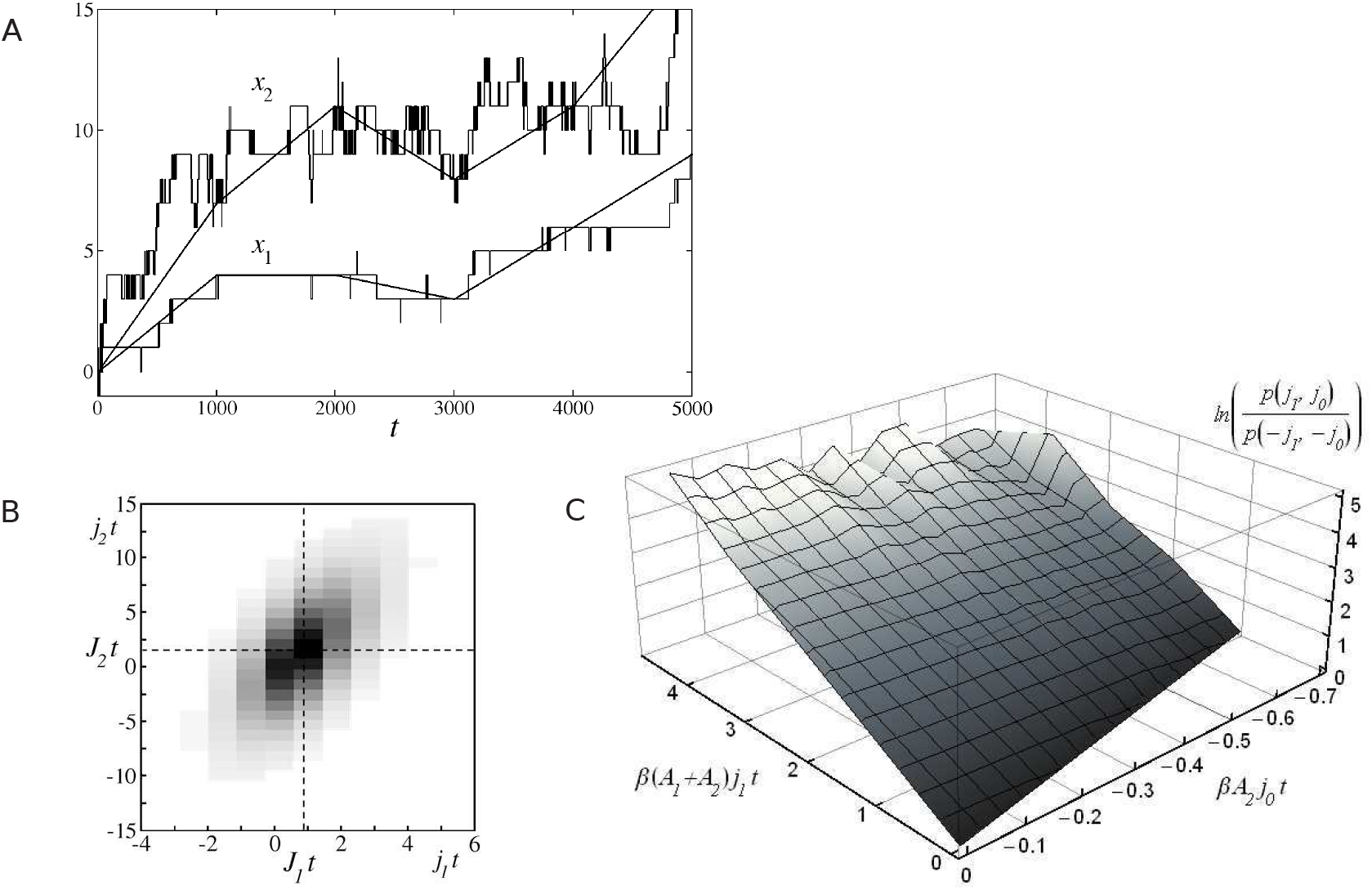
(A) Simulated time course of the net number of external transitions *x*_1_ and *x*_2_ through the input and output gates, respectively, produced in *t* computer steps on the network presented in Fig. 3 E with *βA*_1_ = 1, *βA*_2_ = −0.05 and the fourfold output gate. Determination of the corresponding fluctuating fluxes *j*_1_ and *j*_2_ by averaging over successive time periods of *t* = 1000 steps is shown. (B) The determined Gaussian distribution of the fluxes *j*_1_ and *j*_2_. (C) Numerical verification of the detailed fluctuation theorem 5. Deviations from the plane for large values of *j*_1_*t* and *j*_0_*t* result from insufficient statistical material collected in simulations.

The distribution of random fluxes *j*_1_(*t*) and *j*_2_(*t*) found for *βA*_1_ = 1, *βA*_2_ = −0.05 and the fourfold output gate is depicted in Fig. 5 B. The distribution is actually Gaussian. The thermodynamic behavior of a single biological nanomachine is simple diffusion in a two-dimensional parabolic potential on the plane (*j*_1_, *j*_2_) with the minimum at point (*J*_1_, *J*_2_). After exchanging the variables from *j*_1_ and *j*_2_ to *j*_1_ and *j*_0_, we get the two-dimensional distribution *p*(*j*_1_, *j*_0_). This distribution actually satisfies fluctuation theorem (5), as illustrated in Fig. 5 C.

From two-dimensional distributions *p*(*j*_1_, *j*_0_), we calculated the marginal distributions *p*(*j*_1_) and *p*(*j*_0_) for all simulated trajectories. The logarithms of the ratio of marginals *p*(*j*_i_)*/p*(−*j*_i_) are presented in Figs. 6 A and B as the functions of *j*_i_ *t* for *i* = 1 and 0, respectively. All the dependences are actually linear as predicted by generalized fluctuation theorems 10 and 11.

**Figure 6:**
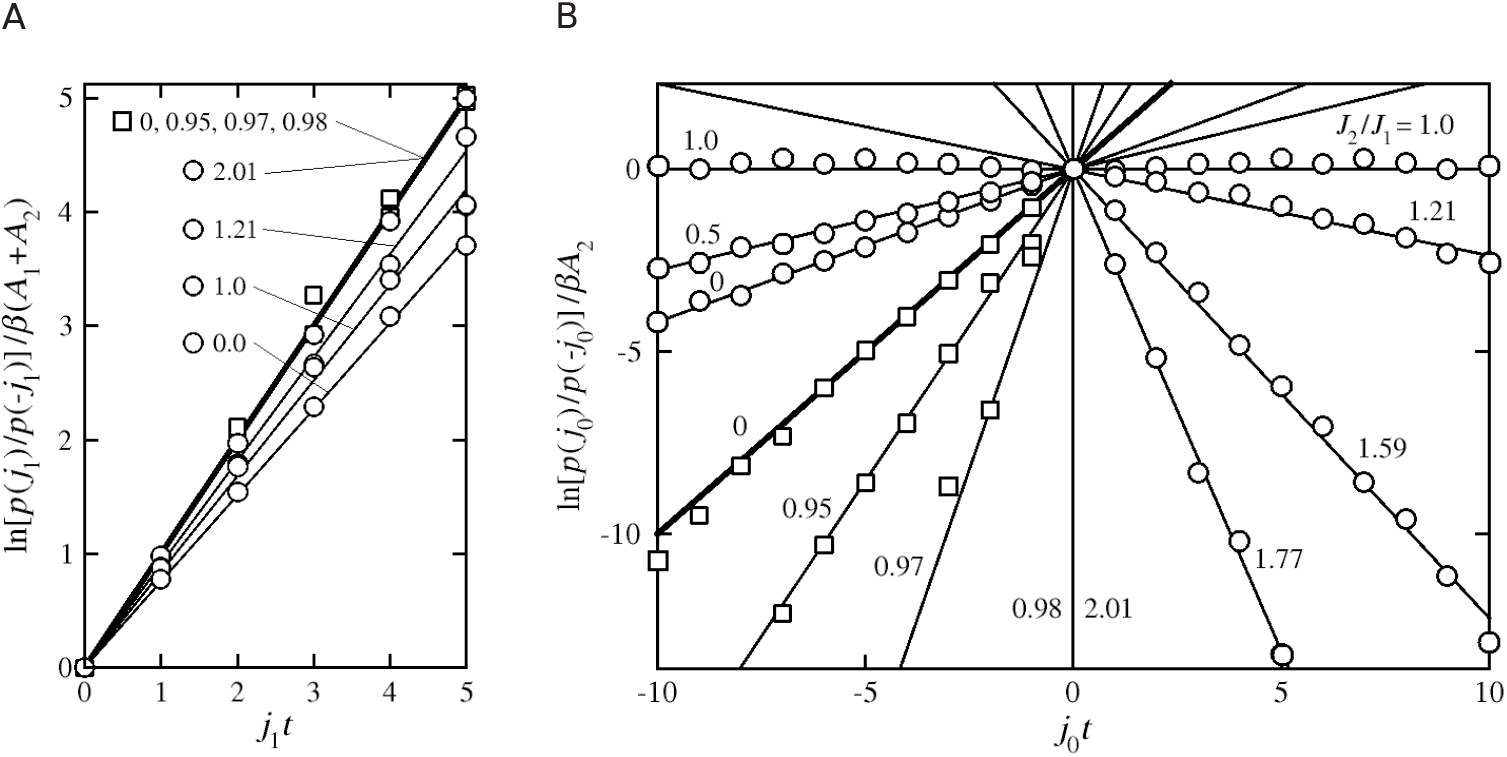
The generalized fluctuation theorem dependences found in the random walk simulations on the network shown in Fig. 3 E with the single output gate (the squares) and the fourfold output gate (the circles). We assumed *βA*_1_ = 1 and a few smaller, negative values of *βA*_2_ determining the ratio of averaged fluxes *ϵ* = *J*_2_*/J*_1_ noted in the graphs. (A) The case of marginal *p*(*j*_1_), compare Eq. (10). (B) The case of marginal *p*(*j*_0_), compare Eq. (11). In order to clearly distinguish the contributions from energy dissipation and information, the results were divided by dimensionless forces *β*(*A*_1_ + *A*_2_) and *βA*_2_ for *i* = 1 and 0, respectively. In this way, the bold lines in both graphs, with unit tangent of the inclination angle, correspond to the dissipation components. Let us recall that *β*(*A*_1_ + *A*_2_) is positive while *βA*_2_ is negative.

For the single output gate (the squares in Fig. 6), we got both *ι*_1_ and *ι*_0_ negative, adding up to positive *σ*_1_ and *σ*_0_. This is consistent with the determined negative values of correlation coefficient *ρ*. Both information contributions differ from zero only for *ϵ* close to unity, i.e. for very small values of force *A*_2_, which does not break practically the detailed balance condition for the transitions through the output gate and is almost a usual white noise. In consequence, information *ι*_1_ is unnoticeable in Fig. 6 A (*A*_1_ + *A*_2_ ≈ *A*_1_), and the information *ι*_0_ in Fig. 6 B, determined mainly by the flux through the single output gate, becomes visible only after dividing by a small value of *A*_2_.

For the fourfold output gate (the circles in Fig. 6), the situation is much more interesting. Now, information contributions *ι*_i_ are of the same sign as dissipation contributions *σ*_i_ and substract from them, which is consistent with the positive values of correlation coefficient *ρ*. The averaged energy dissipations *βD*_i_ = 〈*σ*_i_〉 and informations *I*_i_ = 〈*ι*_i_〉 for *i* = 1 and 0, corresponding to the inclination of the straight lines in Fig. 6 B, are presented in Fig. 7 as the functions of output force *βA*_2_ determining the value of the degree of coupling *ϵ* = *J*_2_*/J*_1_. It is clearly seen that, according to the generalized second law of thermodynamics 15, information *I*_i_ is always less than dissipation *βD*_i_. For *J*_2_*/J*_1_ < 1, which corresponds to large values of output force *βA*_2_, information *I*_0_ is positive, i.e. taken from the microdynamics, to be next subtracted from the positive dissipation *βD*_0_. For *J*_2_*/J*_1_ > 1, which corresponds to small values of output force *βA*_2_, information *I*_0_ is negative, i.e. sent to the microdynamics, and dissipation *βD*_0_ becomes negative. So we showed that the organizational subsystem in our model actually behaves like Maxwell’s demon.

**Figure 7:**
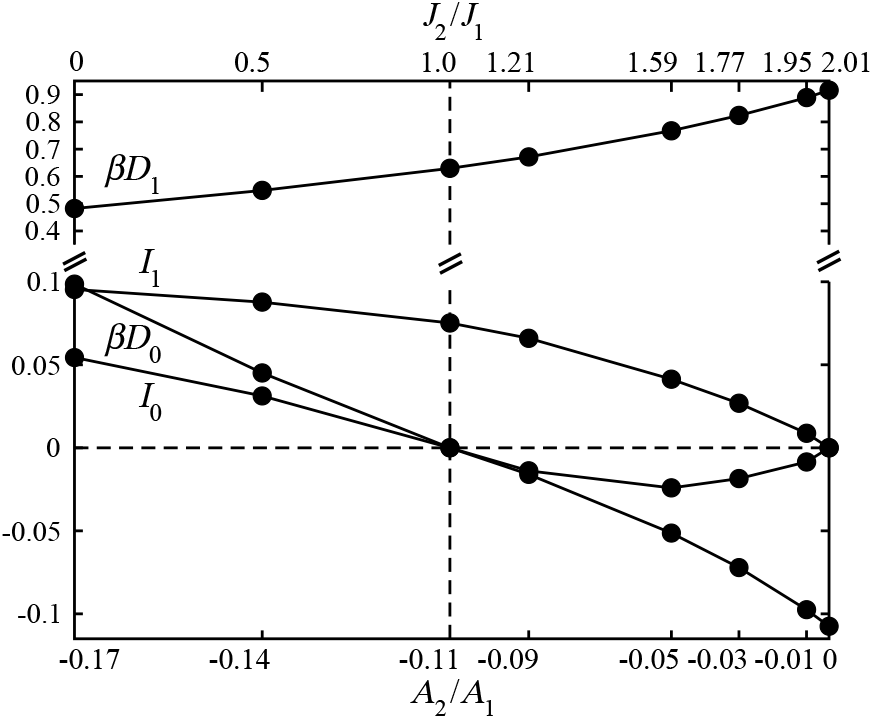
Contribution of dissipation *βD*_i_ and information *I*_i_ to observed entropy production for the studied model system with the fourfold output gate. *i* = 1 in the case of operation energy processing and *i* = 0 in the case of organization energy processing, compare Fig. 4 B. All quantities are presented as function of the ratio of forces *A*_2_*/A*_1_ or fluxes *J*_2_*/J*_1_ and counted in nats (natural logarithm is used instead of binary logarithm) per computer step.

### 3.3 In dimers, biological molecular machines can exchange information with each other

In Fig. 7, which presents the most important result of the paper, the case of tight coupling, *ϵ* = *J*_2_*/J*_1_ = 1, i.e. *J*_0_ = 0 is clearly seen, for which the zero free energy dissipation *D*_0_ occurs simultaneously with the zero transfer of information *I*_0_. The necessary condition for it is a special, critical value of output force *A*_2_ distinguished in Fig. 7 A by the vertical dashed line. There are some arguments that, effectively, such the force value is realized for processive motors with a feedback control between their two identical components [54, 55]. The information transfer to memory takes place for the force of a value less than critical. Only one of the monomers is affected by this force, but the other one, indifferent to external interactions, observes the first monomer and, depending on the result of the observation, exerts an appropriate additional force on it, so that the resulting force reaches the critical value. In consequence, the chosen monomer works as a tightly coupled perfect machine. The process is repeated with the conversion of one monomer into another. The suggested mechanism seems to be in harmony with the consensus kinesin-1 chemomechanical cycle [56] and, in a broader perspective, it could help answer the certainly interesting question: why do most protein machines operate as dimers or higher organized assemblies?

Let us try to formulate this question in a little more detail. In the current model, we did not include intermediate states in the external transitions, replacing them with one computer step. Taking these states into account and giving real value to the times of external transitions requires the replacement of the scheme in Fig. 3 C with the more complex scheme shown in Fig. 8 A. Dynamics in networks M_1_ and M_2_ describe the details of spatial diffusion and molecular recognition in the processes of reactants binding and detaching [39, 47]. Such a model requires the introduction of many new parameters. However, it can be simplified, replacing each transition through the gate in the scheme in Fig. 3 C with the transition to an effective “waiting state” with a long lifetime determined by a fluctuating rate coefficient, as shown in the scheme in Fig. 8 B. We have assumed that the fast transition process under the influence of external force occurs first and the slow recognition process next, but the order can be arbitrary.

**Figure 8:**
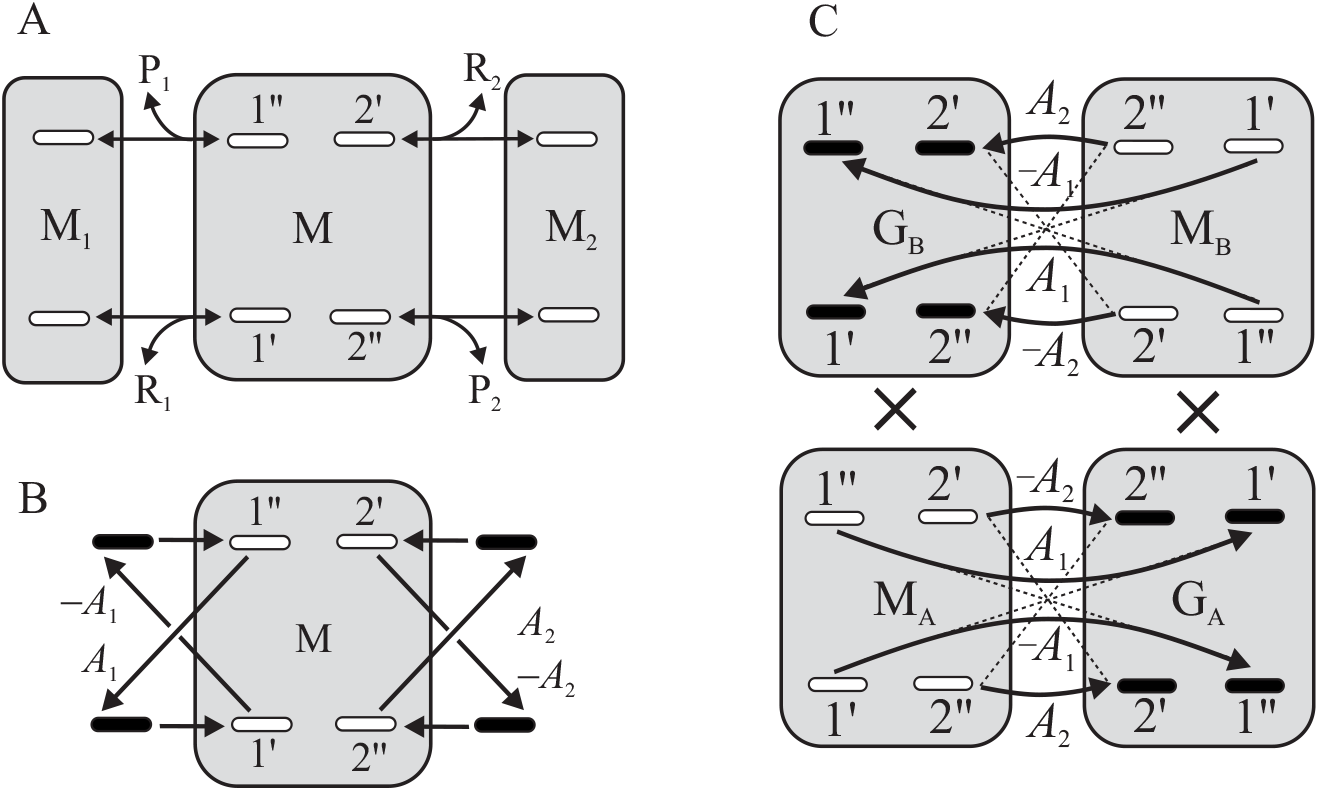
(A) Full scheme of machine’s dynamics in the case where the detaching of products precedes binding of substrates. (B) An effective scheme of dynamics replacing the more detailed scheme above. Transitions to waiting substates, marked as blackened ovals, occur under the influence of external forces, and the further much slower transitions are determined by fluctuating rate coefficients. (C) In dimer, the dynamics of the waiting process in monomer A is imitated by the dynamics of monomer B and vice versa, which is symbolized by the Cartesian product ×. One monomer passes the gate only when the other is frozen in the proper state. When the gate has been passed, its final substate is frozen and the second monomer begins the wander from its own previously frozen substate. The forward transition directions were chosen so that the frozen states of each monomer were in the same half of the doubled network. Changing the frozen states to the wander states is indicated by dashed lines. The gate dimer substates are pairs of gate substates of both constituent monomers, one frozen and the other deciding on the transition, and all dimer dynamics have a bipartite structure.

In the case of dimer, the dynamics of the waiting process in one monomer can be imitated by the dynamics of the second monomer which creates favorable conditions for molecular recognition of reagents. When one monomer is frozen in a certain gate in a state of waiting, the other wanders in its state space and vice versa. In mathematical terms, the states space of the whole dimer has the form of the sum (M_A_ × G_B_) ∪ (M_B_ × G_A_), where indexes A and B label individual monomers. It should be stressed that the space of dimer states is a Cartesian product only locally that means, from a mathematical point of view, it is a fiber bundle.

It is physically reasonable to assume that the monomers are able to recognize their own conformations defined as the regions of influence of the distinguished hubs. Each initial or final gate substate belongs to a specific region of influence. The dynamics of a given monomer is determined by the frozen gate substate of the other. Arriving at its gate substate, the monomer behaves like an autonomous Maxwell’s demon [26, 27, 28, 29, 30, 31, 32]: observes the frozen substate of the second (the “measurement”) and, depending on the energetic advantage, realizes the transition or no (the “feedback control”). When the gate has been passed, its final substate is frozen and the second monomer begins the wander from its own previously frozen substate. The whole situation is illustrated in the scheme presented in Fig. 8 C.

In our simple model from Fig. 3 E, there are only two hubs and, assuming that for the component monomers it is energetically advantageous to be in the opposite conformations, transitions through arbitrary gate are realized only when both the observing and observed gate substates are in the same conformation. Unfortunately, the result of such a procedure turns out to be trivial and nonphysical: we get *J*_2_ = *J*_1_, that means the zero value of *D*_0_ and *I*_0_ in the whole range of values of *A*_2_, but the transitions through the input gate are realized by one monomer, and the transitions through the output gate by the other.

To obtain a non-trivial result for a specific dimer, a specific network with the appropriate number of hubs should be considered. The stochastic thermodynamics of such a nanoscopic system is determined by the dimer gate states, each of which is a pair of gate states of both component monomers, one frozen and one deciding on the transition. This network has a bipartite structure, so our theory determines also the corresponding thermodynamics. Despite the complex formalism, we expect this topic to be worth detailed future research.

## 4 Conclusion

Under physiological conditions, the protein molecular machines fluctuate constantly between lots of conformational substates composing their native state. The probabilities of visiting individual substates are far from the equilibrium and determined by the concentration of the surrounding molecules involved in the process. During the full cycle of free energy-transduction, the possibility to choose different realizations of the free energy-accepting reaction results in the transient limitation of the dynamics to different regions of the conformational network, that is, to breaking the ergodicity [9, 53]. The transient ergodicity breaking makes the machine’s internal dynamics to be a memory for storing and manipulating information. Information is erased each time the energy-donating reaction starts the next cycle. The storage capacity of memory is higher the larger and more complex is the network of conformational substates. This network is particularly large in the case of protein motors, since it then contains the substates of the motor bound to the whole track on which it moves [39], compare Figs. 1 C and 3 B.

Information is exchanged between memory and the thermodynamic variables. Two components of free energy determine the thermodynamic state of the machine. The first, proportional to the concentration of the input reaction product molecules *X*_1_, we refer to as the operation energy. And the second, proportional to the difference in the concentrations of the output and input reaction product molecules *X*_2_ − *X*_1_, we refer to as the organization energy. Similarly as work and dissipation, information is also a change in these well defined functions of state.

If the nanomachine has no choice possibility, both free energy dissipation *D*_1_ and information *I*_1_, as well as *D*_0_ and *I*_0_, are of opposite signs and information processing results in an additional energy loss. Only in the nanomachines that choose randomly the way of doing work and transmit information about it to microdynamics, *D*_0_ and *I*_0_ can be negative, which makes their organizational subsystem Maxwell’s demon. However, according to the fluctuation theorems 10 and 11, the resultant entropy productions in both subsystems must always be non-negative.

At the end, let us take the liberty for a few general remarks. The generalized second laws of thermodynamics 15 were justified only for isothermal processes in the nanoscopic machines. Their universality remains an open problem. One thing is certain. Three necessary conditions must be met to obtain Eqs. 15. These are the presence of an intermediate level of stochastic dynamics, openness providing an external source of free energy, and the appearance of a fluctuating variable characterizing its organization.

Two examples of systems having such properties are well known. The first is thermodynamic systems occurring in the conditions of two phases coexistence, whose organization is specified by an extra thermodynamic variable that survived stochastization [1], referred to as the order parameter [38] or, in various contexts, the emergent [57] or structural [39] variable. In a small enough volume element, this variable is random. An impressive example is the formation of clouds in the conditions of saturated water vapor and air movement. The second example is quantum systems occurring in states entangled with the environment, whose organization is determined by an extra variable that survived decoherence, identified with the classical variable [58, 59]. Here, the value of this variable found in the local measurement with an energy-providing apparatus is random. It is worth investigating whether the stochastic dynamics describing both cases may have a structure that leads to negative free energy dissipation.

## Author Contributions

The general concept and the theory is mainly due to M. K., who also wrote the manuscript. The specification of the critical branching tree model and the numerical simulations are mainly due to P. C.

## Acknowledgements

M. K. thanks Yaşar Demirel and Hervé Cailleau for discussing the problem in the early stages of the investigation.

